# Direct encapsulation of biomolecules in semi-permeable microcapsules produced with double-emulsions

**DOI:** 10.1101/2022.06.23.497329

**Authors:** Grégoire Michielin, Sebastian J. Maerkl

**Affiliations:** Institute of Bioengineering, School of Engineering, École Polytechnique Fédérale de Lausanne

## Abstract

Compartmentalization can serve different purposes such as the protection of biological active substances from the environment, or the creation of a unique combination of biomolecules for diagnostic, therapeutic, or other bioengineering applications. We present a method for direct encapsulation of molecules in biocompatible and semi-permeable microcapsules made from low-molecular weight poly(ethylene glycol) diacrylate (PEG-DA 258). Microcapsules are produced using a non-planar PDMS microfluidic chip allowing for one-step production of water-in-PEG-DA 258-in-water double-emulsions, which are polymerized with UV light into a poly-PEG-DA 258 shell. Semi-permeable PEG shells are obtained by adding an inert solvent to the PEG-diacrylate. Due to the favourable hydrophilicity of poly-PEG-DA 258, proteins didn’t adsorb to the capsule shell, and we demonstrate the direct encapsulation of enzymes, which can also be dried in the capsules to preserve activity. Finally, we leverage capsule permeability for the implementation of a two-layer communication cascade using compartmentalized DNA strand displacement reactions. This work presents the direct encapsulation of active biomolecules in semi-permeable microcapsules, and we expect our platform to facilitate the development of artificial cells and generating encapsulated diagnostics or therapeutics.

## Introduction

Recent advances in microfluidics, material science, synthetic biology, and bioengineering enable precision engineering of micron sized compartments, allowing the manipulation of increasingly complex biomolecular systems[1, 2]. Droplet microfluidic technology enables the production of polymeric compartments with precisely defined size and structure, templated from double-emulsions or higher order emulsions with a polymerizable phase as reviewed by Datta *et al*.[3]. One drawback with polymer precursors used as a middle-phase is that the obtained polymerized compartments are in general hermetic to the permeation of solutes, however Kim *et al*.[4] demonstrated that semi-permeable polymeric shells can be fabricated by introducing an inert diluent, or porogen, in the polymer precursor phase. Upon UV polymerization, the non-reactive porogen is excluded from the polymerizing polymer in a process called polymerization-induced phase separation (PIPS). However, the drawback of the polymers used in this study and of most polymers obtained from water-immiscible precursors is the hydrophobicity of the polymerized capsule shell leading to protein adsorption, and therefore preventing direct encapsulation of proteins or enzymes.

Recently it was shown that PEG-diacrylate with a molecular weight of 258 g/mol PEG-DA 258 can be used as a polymerizable middle phase for the production of double-emulsions with a microfluidic capillary device[5]. The researchers also showed that poly-PEG-DA 258 has a hydrophilicity similar to hydrogels made from water soluble PEG-DA derivatives of higher molecular weight. Such favorable characteristics allowed them to directly encapsulate fibrinogen without noticeable adsorption to the polymer shell. However, the produced capsules were also hermetic and prevented even small molecule dyes (erioglaucine disodium salt: 793 g/mol, oil red O: 408 g/mol) to diffuse out of the capsules, which would severely limit the use and application range of such capsules.

To directly encapsulate biologically active macromolecules in biocompatible semi-permeable capsules, we combined the favorable characteristics of the low molecular weight PEG-DA 258 with pore formation by PIPS. In this study, we used a PDMS microfluidic device with a flow-focusing, co-axial, and non-planar geometry which does not require surface treatment to produce doubleemulsions of water-in-PEG-DA 258-in-water. In order to obtain semi-permeable capsules, we used PIPS to form small pores in the capsule shells. By encapsulating fluorescent cargoes of different sizes or placing empty capsules in fluororophore containing solutions, we showed that the capsules were semi-permeable with a size cutoff allowing the direct encapsulation of proteins and enzymes of 32.7 kDa and above, while permitting transport of smaller molecules. Biomolecules were retained in the capsules without degradation, and were homogeneously distributed inside the capsules. Encapsulated enzymes survive air drying in trehalose and had enzymatic activity after rehydration. The semi-permeable capsules were able to communicate with their environment allowing us to implement a two-layer signalling cascade by immobilizing DNA strand displacement reactions inside the liquid core of two different microcapsule populations[6]. Altogether, this work describes the development and characterization of semi-permeable microcapsules with a biocompatible polymeric shell that can be used for the encapsulation of different biomolecules for use in diagnostic, therapeutic, or synthetic biology applications.

## Results

### Production of semi-permeable PEG-DA 258 microcapsules using PIPS

As PEG derivatives are generally considered biocompatible, bioinert, and biodegradable[7], we tested various PEG-DAs of low molecular weight, such as PEG-DA MW 700, PEG-DA MW 575 and PEG-DA MW 258 as precursors for the production of microcapsules after photopolymerization by UV illumination. For PEG-DA with MW 575 or higher, their water miscibility prevented us to readily form double-emulsions. While Leonavicene *et al*. recently showed that it is possible to obtain a two-phase system by combining PEG-DA MW 575 and high molecular weight PEG-DA MW 8000 in the inner phase and form core-shell capsules using PEG-DA as a capsule material[8]. However, we considered an alternative approach in using PEG-DA MW 258 as a potential polymer precursor due to its water immiscibility, as was also reported by Nam *et al*.[5]. This property of PEG-DA MW 258 allowed us to use it as the middle phase in double-emulsions and was compatible with droplet generation in a PDMS device (Figure 1A). To form the double-emulsions, we used an aqueous continuous phase supplemented with 10% PVA. The PEG-DA 258 middle phase is supplemented with a photoinitiator (HMPP) and surfactant (Span80), with the optional addition of a mild organic solvent. We produced monodisperse double-emulsions in a jetting regime with flow rates of 2500 *µ*L/hr for the aqueous continuous phase, 200 *µ*L/hr for the PEG-DA MW 258 middle phase, and 250 *µ*L/hr for the aqueous inner phase (Figure 1, B). Interestingly, the doubleemulsions could be generated without requiring any surface treatment of the device owing to the non-planar geometry of the PDMS device which prevents wetting of the collection channel by the hydrophobic middle-phase.

**Figure 1:**
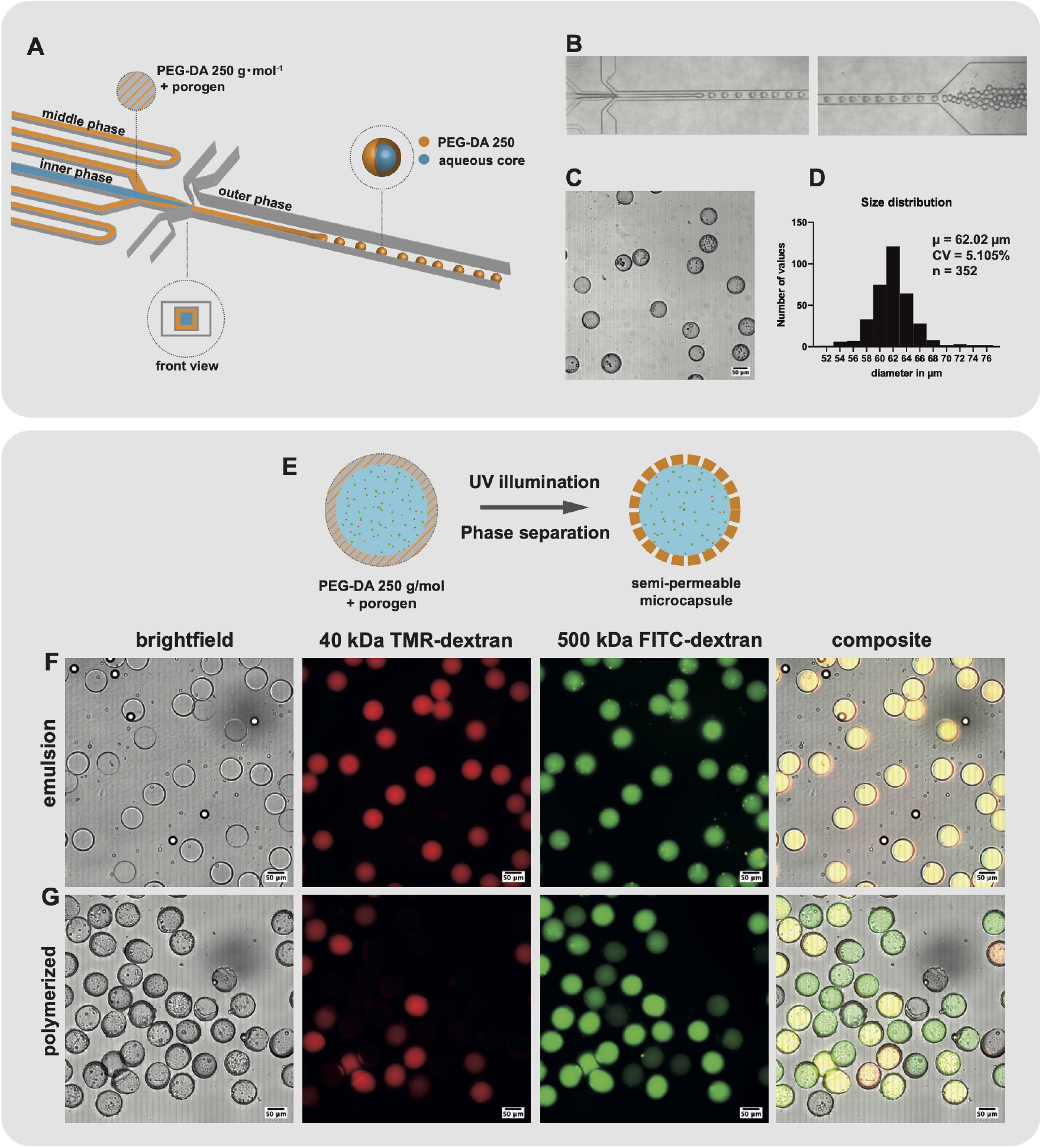
Production of semipermeable microcapsules in a PDMS device with 3D geometry. **(A)** Schematic representation of the PDMS device with 3D geometry. W/O/W doubleemulsions are generated with a PEG-DA 258 middle phase encapsulating an inner aqueous core. **(B)** Micrographs of the PDMS device operation. **(C)** Brightfield image of microcapsules obtained after UV polymerization of the collected double-emulsion. **(D)** Size distribution of a representative batch of polymerized microcapsules. **(E)** Schematic representation of PIPS upon UV illumination. 15% Butyl-acetate (porogen) was mixed with PEG-DA 258 to form semi-permeable microcapsules. **(F)** In the collected double-emulsion, both high molecular weight 500 kDa FITC-dextran and lower molecular weight 40 kDa RITC-dextran are retained in the inner aqueous phase. **(G)** After UV polymerization and PIPS, pores are formed in the capsule shell and capsules become semipermeable.

We collected double-emulsions in a UV-transparent cuvette for 15 minutes, after which the capsules were polymerized in batch by UV exposition. After polymerization we obtained monodisperse capsules with a mean diameter of 62 *µ*m (Figure 1C,D). The coefficient of variation was close to 5%, in accordance with previous results using similar PDMS devices[9]. The capsule shell was estimated from inspecting microscope images to be between 5 and 10 *µ*m thick. Our results not only confirm the observation from Nam *et al*.[5] that water-immiscible PEG-DA MW 258 is a suitable middle phase, but demonstrated its compatibility with double-emulsion generation in a PDMS device which does not require any surface treatment, and resulted in the production of monodispersed poly-PEG-DA 258 microcapsules with porous thin shells (Figure 1E-G).

To produce semi-permeable capsules, we used the mild inert solvent butyl-acetate added to the middle phase serving as a porogen in PIPS[10, 4]. Butyl-acetate was used by Kim *et al*.[4] to form nanopores with diameter below 30 nm in a thin shell composed of a cross-linked network of ethoxylated trimethylolpropane triacrylate (ETPTA) and glycidyl methacrylate (GMA). To evaluate the permeability of the semi-permeable capsules, we added 500 kDa FITC-dextran and 40 kDa TMR-dextran to the inner aqueous phase, and observe that almost 90% of the polymerized capsules retained the 500 kDa FITC-dextran after the capsules were extensively washed by successive centrifugation in aqueous buffer and followed by a 24h incubation in buffer at 4^°^C (Figure 1G). On the other hand, only about 40% of the capsules retained the lower molecular weight 40 kDa TMR-dextran. These results showed that some capsules are semi-permeable, as evidenced by capsules in which only the higher molecular weight fluorophore is retained. Even though the size cut-off could not be precisely determined from this experiment, we estimated it to be well below 500 kDa and potentially close to the size of the smaller fluorophore of 40 kDa.

To better characterize the semi-permeability, we prepared empty capsules and placed them in a solution of fluorescent molecules of smaller molecular weights (Figure 2). We placed empty capsules produced using 15% butyl-acetate porogen in a solution containing both 10 kDa RITC-dextran and 32.7 kDa EGFP (Figure 2A,B). We observed that most capsule showed a relatively rapid increase in signal in the red fluorescent channel, corresponding to the lower molecular weight 10 kDa RITCdextran. After 1 hour incubation, most capsules showed a red fluorescent signal, and all capsules displayed a red fluorescent signal after 24h incubation. At the same time, the majority of these capsules showed no signal increase in the green fluorescent channel corresponding to the 32.7 kDa EGFP. By superimposing the signal from the two fluorescent channels, we could clearly see that a majority of the capsules were semi-permeable and excluded EGFP for at least 24h while allowing diffusion of the 10 kDa RITC-dextran into their interior. We observed however some variability in the permeability of the capsules, with some capsules appearing permeable to EGFP already after a few minutes of incubation, while a few capsules still excluded 10 kDa RITC-dextran after 1 hour. We obtained semi-permeable capsules with a different size cutoff using 1-decanol as a porogen.

**Figure 2:**
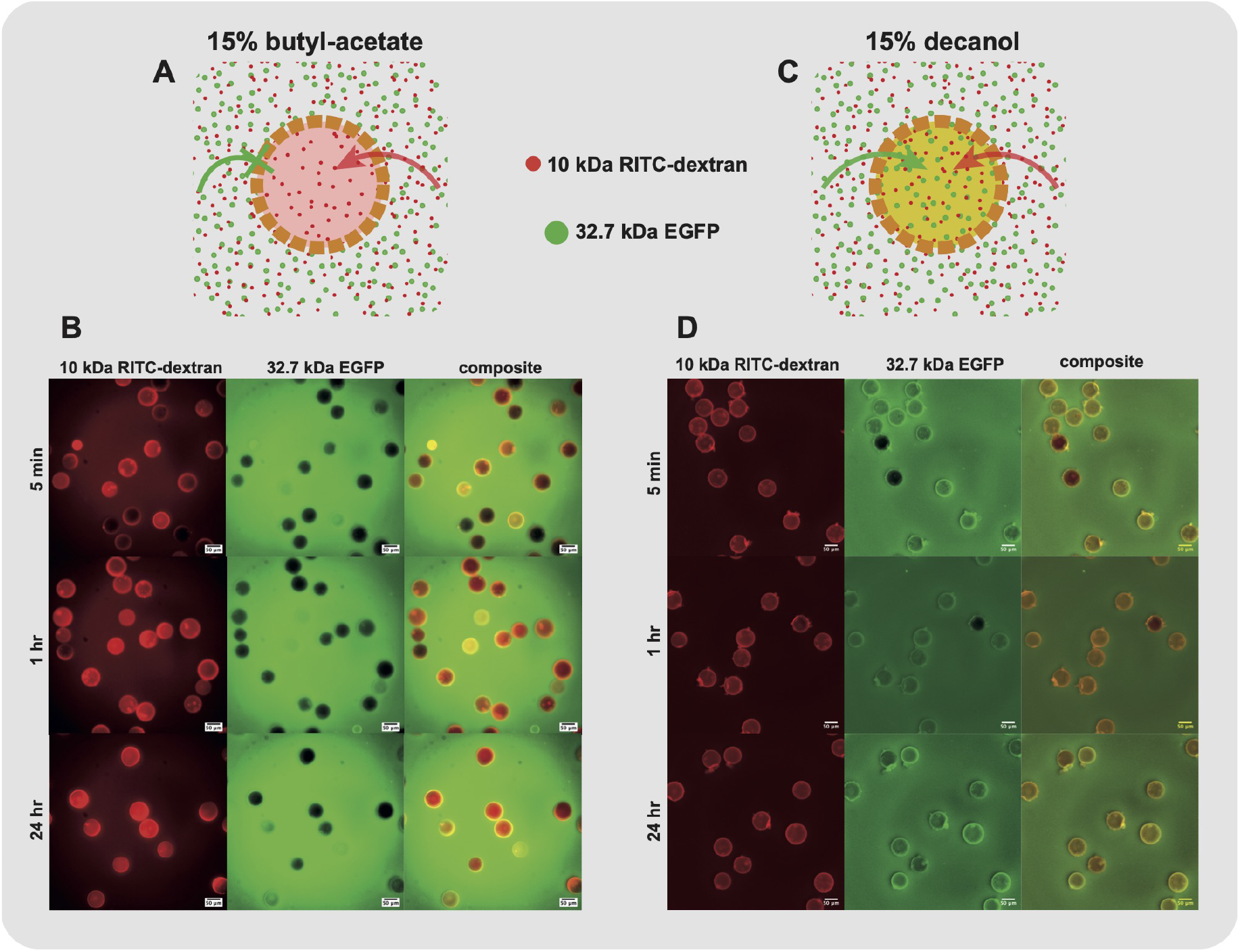
PEG-DA 258 microcapsules are semi-permeable and the pore size can be adjusted by changing the porogen. Schematic representation of PEG-DA 258 microcapsules produced using **(A)** 15% butyl-acetate or **(C)** 15% 1-decanol as porogen. 15% butyl-acetate microcapsules selectively allowed the permeation of 10 kDa RITC-dextran while excluding 32.7 kDa EGFP. The larger pore size of microcapsules produced with 15% 1-decanol as porogen allowed the diffusion of both fluorescent molecules. Microcapsules produced using **(B)** 15% butyl-acetate or **(D)** 15% 1-decanol porogen were immersed in a solution containing 10 kDa RITC-dextran and 32.7 kDa EGFP. The evolution of the fluorescent signal in the Cy3 and FITC channels was observed after 5 minutes, 1 hour and 24 hours. While microcapsules produced using 15% 1-decanol were permeable to both fluorescent molecules, we clearly observed the selective permeability of microcapsules produced using 15% butyl-acetate as a porogen.

The use of 1-decanol was reported by Kim *et al*.[4] to create larger pores in the ETPTA/GMA polymer shell due to a different interaction parameter of 1-decanol with the forming polymer. Here, we show that capsules produced with 15% 1-decanol porogen in PEG-DA 258 also resulted in higher permeability than capsules produced with the butyl-acetate porogen (Figure 2C,D). We observed an increase in green fluorescent signal corresponding to 32.7 kDa EGFP in the capsule interior after only a few minutes incubation and all capsules were fluorescent after 24 h. In addition, all capsules were completely permeable to 10 kDa RITC-dextran. These results demonstrate that it is possible to tune microcapsule permeability by varying porogen composition, and that butyl-acetate and 1-decanol are compatible with our microfluidic production of semi-permeable microcapsules.

### Direct encapsulation of proteins

Next, we show that we can directly encapsulate proteins in semi-permeable poly-(PEG-DA 258) microcapsules without adsorption to the polymeric shell or loss of function. We encapsulated 32.7 kDa EGFP inside our microcapsules by adding it to a final concentration of 2 *µ*g/mL to the inner aqueous phase. EGFP fluorescence signal was observed in the double-emulsion without any sign of precipitation, and, once polymerized, the capsules displayed a homogeneous fluorescent profile without any sign of protein adsorption to the capsule material (Figure 3E,G). We also encapsulated 60 kDa FITC-labelled streptavidin in the aqueous inner phase to a final concentration of 50 *µ*g/mL, and the fluorescent signal profile in the polymerized capsules indicated no sign of protein adsorption on the capsule shell (Figure 3I,K). After incubation in PBS for 24h, we saw that a proportion of the EGFP-containing capsules did not contain any fluorescent signal (Figure 3e,f). This observation suggests a permeability cutoff close to the size of these fluorescent biomolecules. We also expect some variability in the pore size of the capsules due to the batch UV polymerization process, during which the UV intensity could slightly differ depending on the position of the capsule in the cuvette.

**Figure 3:**
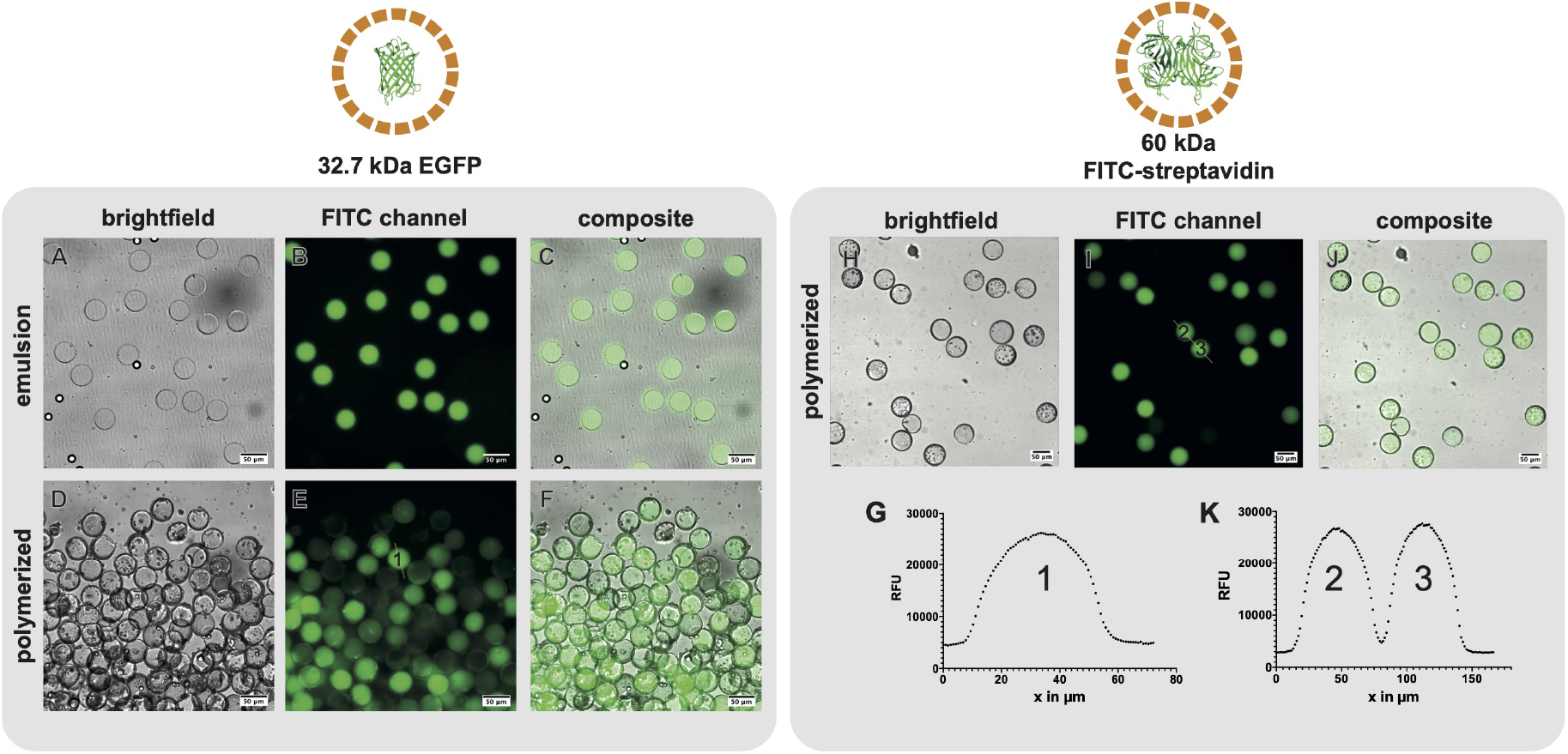
Direct encapsulation of proteins inside semi-permeable PEG-DA 258 capsules. EGFP was added to the aqueous inner phase for direct encapsulation. **(A, B, C)** Microscope images of double-emulsions showing a fluorescent signal in the FITC channel. **(D, E, F)** After polymerization, the fluorescent signal was still present in the interior of most capsules. Variability in the pore size or a size cutoff close to the 32.7 kDa EGFP resulted in some protein leakage. **(G)** Fluorescent intensity profile of the capsule indicated in panel E. The fluorescent profile suggests a homogeneous distribution of EGFP in the interior of the capsule without protein adsorption to the shell material. **(H, I, J)** Microscope images of polymerized capsules containing FITC-streptavidin after direct encapsulation. Fluorescent signal is present in all capsules, suggesting that the pore size is too small for FITC-streptavidin leakage. **(K)** Fluorescent profile across two capsules from panel I. The profile suggests a homogeneous distribution of FITC-streptavidin in the interior of the capsule without protein adsorption to the shell material.

Also, some capsules might have been broken or damaged which would lead to the release of their cargo. With the larger molecular weight 60 kDa FITC-streptavidin, we saw that most polymerized capsules contained fluorescent signal after aqueous washes, indicating a size cutoff below the size of FITC-streptavidin.

We demonstrated that the poly-PEG-DA 258 shell obtained after UV polymerization is compatible with direct protein encapsulation. The materials used did not lead to adsorption of proteins to the capsule shell and the polymerization-induced phase separation process formed pores sufficiently small to retain proteins with molecular weights of 32.7 kDa and above.

After successful encapsulation of fluorescent proteins, we encapsulated enzymes and performed enzymatic assays with the produced capsules. We used a recombinant GFP-luciferase fusion protein (econoLuciferase™, Biosynth), which allowed confirmation of encapsulation by measuring its fluorescent signal. The molecular weight of the fusion protein being over 90 kDa led to retention of the enzyme in the interior of the microcapsules. Indeed, we saw that both the double-emulsion and polymerized capsules contained a fluorescent signal from the econoLuciferase fluorescent fusion protein. In both cases, we observed a speckled distribution of the fluorescent signal which might be due to some precipitation in the 10% PVA inner solution (Figure 4A,B). The econoLuciferase-containing capsules were placed in a solution containing D-luciferin, and we measured a bioluminescent signal two orders of magnitude higher than the signal observed for empty capsules (Figure 4C). These results demonstrated that an active enzyme can be encapsulated in poly-PEG-DA 258 microcapsules, with the semi-permeable shell allowing diffusion of the substrate D-Luciferin into the core of the microcapsules to generate a bioluminescent signal.

**Figure 4:**
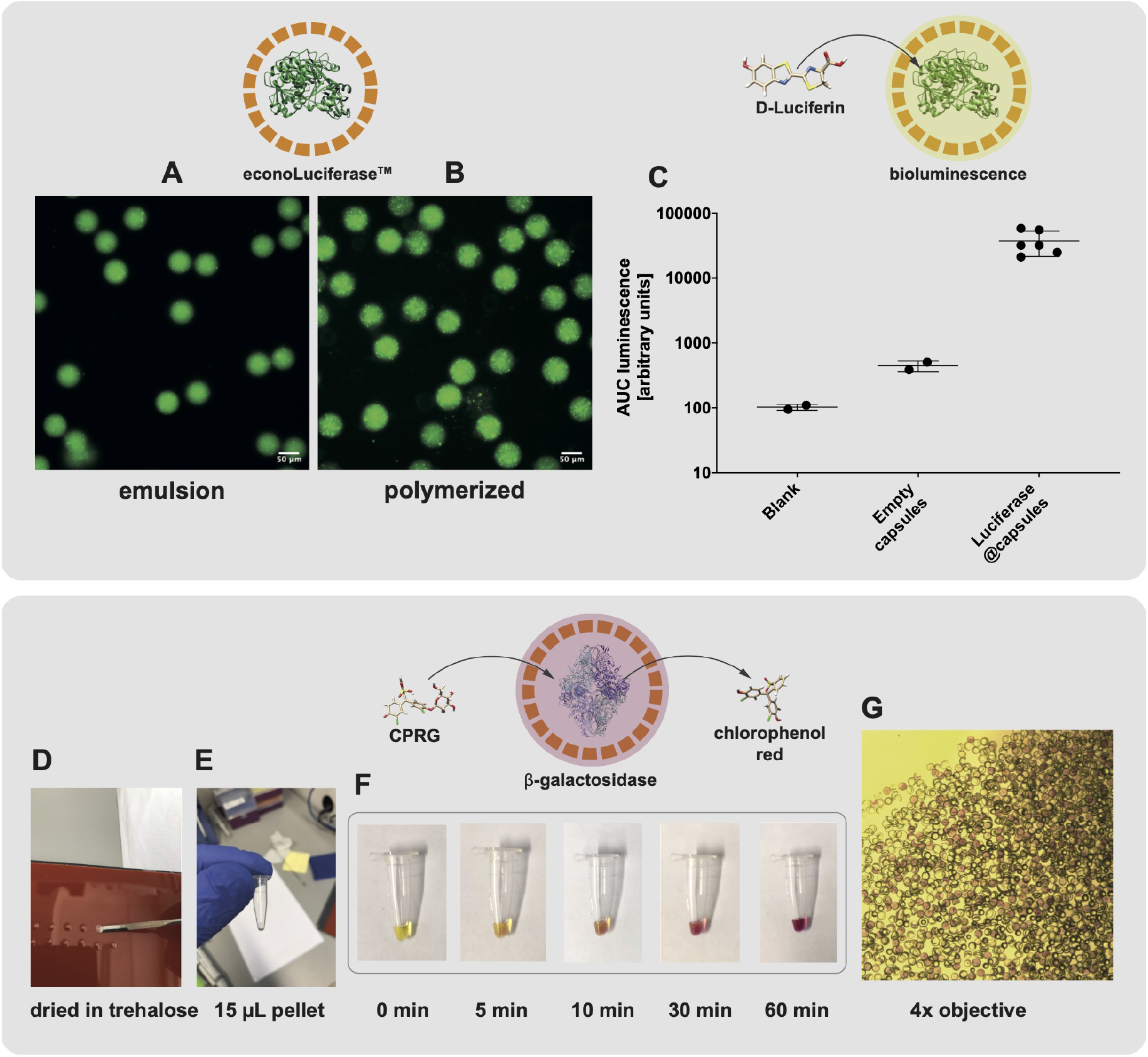
Direct encapsulation of functional enzymes in semi-permeable PEG-DA 258 microcapsules. A luciferase-GFP fusion protein was directly encapsulated in semi-permeable capsules. The fluorescent fusion protein allows for the visualization of the enzyme **(A)** in the double-emulsion, and **(B)**, in the polymerized capsules. **(C)** The encapsulated econoLuciferase™ shows a strong signal in a bioluminescent assay. Direct encapsulation of *β*-galactosidase. **(D, E)** Enzyme-containing capsules were dispersed in trehalose and air-dried at 37^°^C. **(F)** After rehydration with a solution containing CPRG, the substrate was hydrolyzed to chlorophenol red. **(G)** The solution was imaged with a color camera mounted on an inverted microscope with 4x magnification. Capsules displayed a purple color in their interior, indicative of *β*-galactosidase enzymatic activity.

In a second example, we showed that capsules can be dried and rehydrated while preserving enzyme function. We encapsulated *β*-galactosidase, a tetrameric enzyme with a molecular weight of 465 kDa. To show that enzyme-containing capsules can be dried and retain activity upon rehydration, we dispersed *β*-galactosidase-containing capsules in a 0.5 M trehalose solution and dried small drops overnight in an incubator at 37^°^C resulting in trehalose pellets (Figure 4D,E). After rehydrating the dried pellets with a Chlorophenolred-*β*-D-galactopyranoside (CPRG) solution, we observed a change in color upon enzymatic conversion of the yellow CPRG to chlorophenol red. We observed that the color change occurred at the location of the capsules, and visualization with a 4x objective showed that the interior of some caspules turn to an intense purple red color (Figure 4F,G). Although the images were not used as a quantification of the chlorophenol red concentration, they clearly showed that conversion of CPRG to chlorophenol red occurred inside the *β*-galactosidase-containing capsules. These results demonstrated the possibility for the direct encapsulation of active enzymes into semi-permeable microcapsules, and we additionally showed that capsules can be air-dried in a lyoprotective solution of trehalose and still retain their activity after rehydration.

### Encapsulated DSD reaction networks and implementation of a two-layer signalling cascade

We investigated the possibility of encapsulating more complex biomolecular systems, as this could be used for sensing and responding to a stimulus in diagnostic, therapeutic, or theranostic applications. It was recently demonstrated that DNA strand displacement (DSD) reactions can be performed in proteinosome microcompartments[11] as a model of protocellular communication and distributed biomolecular computation[6]. Here, we immobilized biotinylated DNA strands in our poly-(PEG-DA 258) capsules containing streptavidin, following the design from Joesaar *et al*.[6].

We implemented a two-layer signalling cascade by functionalizing a first population of capsules with a transducer DSD gate that activates after toehold displacement by a ssDNA input strand (A), which leads to the release of a signal strand (Q1) and unquenching of a Cy5 fluorophore. The (Q1) signal strand can in turn activate a second population of capsules functionalized with a transducer-amplifier DSD gate releasing a second signal strand (Q2), this time unquenching a Cy3 fluorophore. We also added a fuel strand which acts as an amplifier (Figure 5A). The two capsule populations were mixed together and the two-layer signalling cascade was activated upon addition of the input strand (A). In this experiment, we mixed the two capsule populations, added 50 nM input strand (A), and loaded the freely moving capsules into a cell counting chamber. We first measured a Cy5 signal increase corresponding to the first population of capsules containing the first DSD transducer gate being activated. After a time lag of a few minutes, we saw a subsequent increase in Cy3 signal corresponding to the second population of capsules containing the transducer-amplifier DSD gate (Figure 5B,C). We observed a Cy5 increase in about 10 capsules of the first population and Cy3 increase in about 15 capsules of the second population. The activation of the two-layer cascade could be modified by changing the concentration of input strand (A). By increasing its concentration to 100 nM, the activation of the first population was greatly accelerated, and it was difficult to capture the initial signal increase (Supplementary Figure S1). On the other hand, reducing concentration of (A) to 10 nM led to a much lower level of activation (Supplementary Figure S1). In all cases, we observed a lag of 5 to 10 minutes between activation of the first population and activation of the second population. While this is a succinct implementation of the recently developed compartmentalized DSD reactions, these results demonstrated that DSD reactions can be efficiently encapsulated in our semi-permeable microcapsules and employed to build communicating biomolecular systems.

**Figure 5:**
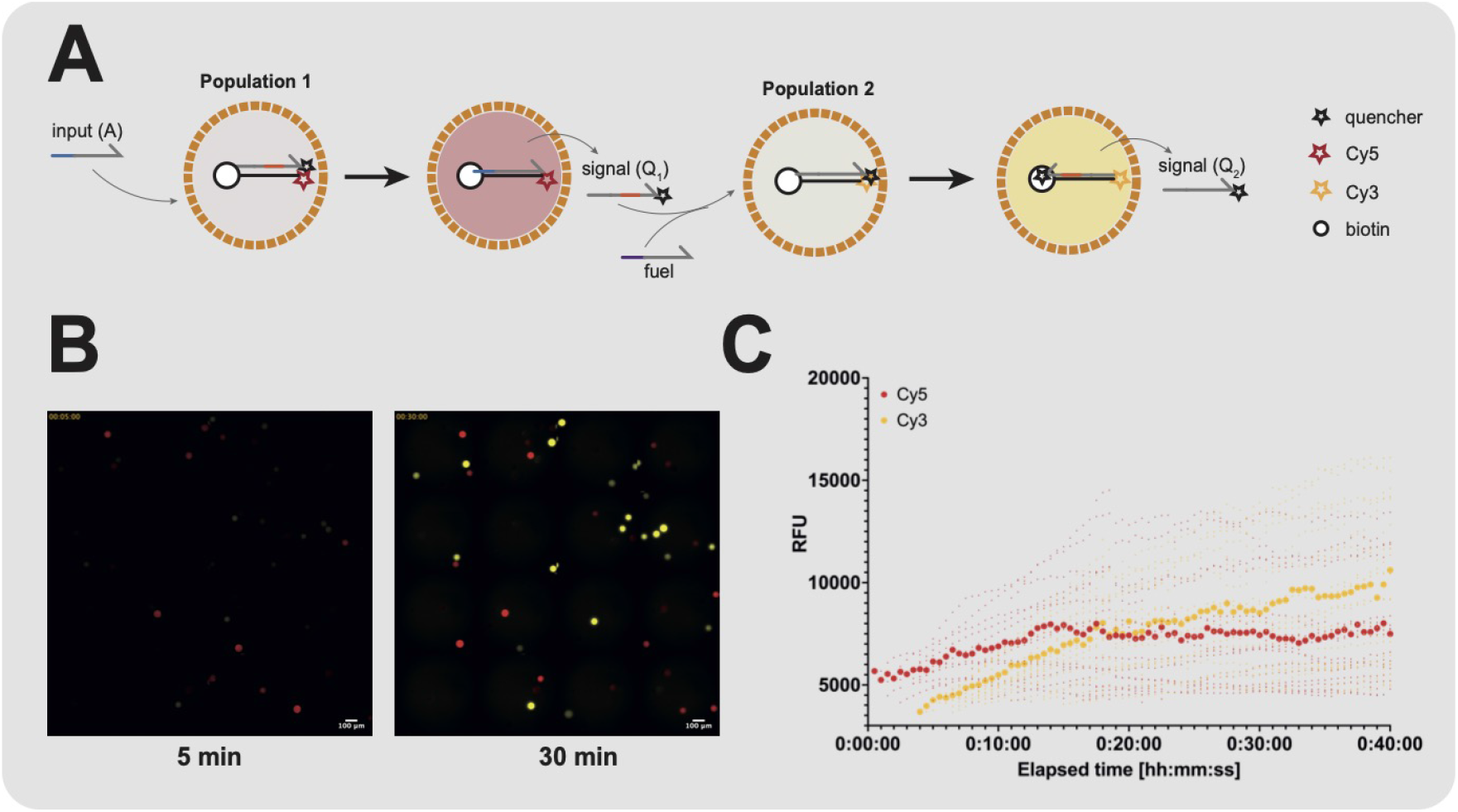
Immobilization of DNA strand displacement reaction network in semipermeable microcapsules and implementation of a two-layer signalling cascade. **(A)** Schematic representation of the two-layer signalling cascade as developed by Joesaar *et al*. [6] **(B)** Implementation of the two-layer signalling cascade in poly-PEG-DA 258 capsules. The two capsule populations were mixed together and imaged on a cell-counting slide immediately after addition of 50 nM input strand (*A*). An increase in Cy5 and Cy3 fluorescent signals was observed corresponding to the activation of the first and second populations, respectively. **(C)** Median intensity of detected particles. An increase in the Cy5 signal was observed corresponding to the activated first population of capsules. After release and diffusion of the signal strand (*Q*_1_) to the second capsule population, an increase in Cy3 signal was observed corresponding to their subsequent activation. The larger symbols correspond to the median of all detected particles in a given fluorescent channel.

## Discussion

In this work, we present the production of biocompatible and semipermeable poly-PEG-DA 258 microcapsules and their use for encapsulation of proteins or DNA networks while retaining functionality. First, we show that PEG-DA 258 can be used as a polymerizable middle phase for the production of microcapsules templated from water-in-PEG-DA 258-in-water double-emulsion. The generation of such double-emulsion does not require the use of a glass capillary device[5], but can be produced in a PDMS device with 3D geometry which does not require any surface treatment[12]. The relatively simple method and reproducibility in the fabrication and use of untreated PDMS devices for the production of poly-PEG-DA 258 microcapsules should make this method available to many labs with access to basic soft lithography fabrication.

By adding an inert diluent to the PEG-DA 258 middle phase, we show that semipermeable microcapsules can be formed by PIPS. This technique was previously used in the formation of sermi-permeable microcapsules made from other acrylate based polymers such as blends of ETPTA/GMA[4] or EDGMA/GMA[10]. While the semi-permeability of capsules made from such polymers was demonstrated, no direct encapsulation of biological components was achieved, apart from the encapsulation of plasmid DNA in the ETPTA/GMA microcapsules in Niederholtmeyer *et al*.[13]. It is interesting to note that the plasmid DNA contained in these capsules could still serve as a protein expression template after immersion in a cell-free transcription translation system. However, the ETPTA/GMA capsule had to be reacted first with amino-PEG12-alcohol to prevent the adsorption of proteins on the capsule shell, which also indicates that direct encapsulation of proteins or enzymes would have been precluded by the use of such polymers.

Here, we confirm previous observation from Nam *et al*.[5] that poly-PEG-DA 258 microcapsules do not lead to the adsorption of biomolecules, including fluorescent proteins or enzymes, allowing their direct encapsulation in the interior of the semi-permeable microcapsules. Moreover, the small pores obtained by PIPS when using butyl-acetate as a porogen were estimated to be close to or even smaller than the hydrodynamic radius of the 32.7 kDa small EGFP. This will allow for the encapsulation of a variety of proteins, RNA, DNA or other (bio)-molecule of interest as long as their size is not smaller than this cut-off. We show that the encapsulation of active enzymes is possible and that the semi-permeability of the capsules overcomes the challenges of stable encapsulation without release of the enzyme, while at the same time allowing diffusion of reactants and products in and out of the capsules. Alternatively, we propose that the encapsulation of streptavidin can serve as an immobilization partner for smaller molecules. Also, encapsulation of functionalized magnetic beads or particles of appropriate size, could be used for the same purpose. In future developments, this could serve to immobilize peptides and small proteins with affinity tags, or even small molecules, which could be activated or released upon sensing of an external stimulus[14]. Here we applied this concept by immobilizing short strands of DNA modified with a biotin handle on encapsulated streptavidin, which served in the implementation of a two-layer signalling cascade from DNA strand displacement (DSD) reactions. While the implementation of DSD reactions in microcompartments proposed by Joesaar *et al*.[6] was originally performed in proteinosomes, we show that our semi-permeable microcapsules are also capable of implementing such complex synthetic biology application.

In conclusion, we demonstrated the one-step encapsulation of a variety of biomolecules in polyPEG-DA 258 microcapsules templated from double-emulsions. Fluorescent molecules, proteins, enzymes, and DNA strands with biological activity can be loaded in the core of the biocompatible semi-permeable capsules. This opens doors towards building complex and robust multi-component biomolecular artificial cells with diagnostic, therapeutic, and synthetic biology applications.

## Acknowledgments

We thank Rohan Thakur for his contribution in setting up the syringe pumps and for his contributions in initial platform development. We thank Jui-Chia Chang, Gianluca Etienne, and the SMaL lab at EPFL, for supplying us with the microfluidic device photomask and for their help with preliminary experiments. GM was supported by the MD-PhD Program of the Swiss National Science Foundation (SNF, 323530 171144) and by a SNF NRP (National Research Program) 78 Covid-19 Grant 198412 (to S.J.M.). This work was published as part of EPFL MD-PhD thesis N°8385[15].

## Author contributions

G.M. and S.J.M. designed research; G.M. performed research; G.M. and S.J.M analyzed results and data; G.M. and S.J.M. wrote the manuscript.

## Supporting Information

Materials and Methods Supplementary Figures and Tables

## Materials and methods

### Microfluidic chip fabrication

The PDMS microfluidic devices were produced using standard soft lithography methods based on designs previously published [1, 2]. Transparency films obtained from the SMaL lab from Prof. Esther Amstad at EPFL were used as photo masks for the production of a SU-8 master on a silicon wafer. A master was prepared using photolithography with the negative photoresist SU-8. A first 40 *µ*m layer of 3025 SU-8 (Microchem) containing the inner and middle phase channel was spincoated and UV exposed on the first half of the master while the second half was fully exposed. A second 60 *µ*m layer of 3050 SU-8 (Microchem) was spin-coated and UV exposed aligned on top of the features to generate the main channels on both halves of the master. After using the master for PDMS replica moulding, both halves of the device were treated with oxygen plasma for 45 s and bonded to one another.

### Microfluidic microcapsules production

The microfluidic system was operated using syringe pumps (New Era) controlled via a python script written in Professor Adam Abate’s lab at UCSF. Syringes were connected to Tygon tubing and the tubing was connected to the chip via metal pins. The flowrates for the inner, middle and outer phase were set to 250, 200, and 2500 *µ*L/h. The inner phase and outer phase are composed of a 10 wt/wt% aqueous solution of PVA (poly(vinyl alcohol), 13000-23000 g/mol, Sigma-Aldrich). Droplet production was observed at the junction and at the collecting channel through a 4x objective with a high-speed digital camera (Fastec HiSpec). The resulting double-emulsions were placed on a cell-counting slide and inspected with a 20x objective in brightfield and fluorescence. The double-emulsions were collected in UV-transparent cuvettes (UVettes, Eppendorf) and illuminated for 30 seconds from the side with the complete spectrum of a 100W Mercury lamp focused on the cuvette. Alternatively, an Omnicure S1500 200W UV curing lamp with standard filter (320nm-500nm) was used and the double-emulsions were illuminated with a probe placed on the top of the cuvette. The polymerized capsules were transferred to a 1.5mL Eppendorf tube and washed 10 times in 1 mL cold wash buffer (1xPBS, 0.1 % Tween20) by centrifugation at 2000 rpm for 1.5 minutes at 4^°^C. Capsules were stored at 4^°^C in wash buffer.

### Semi-permeable capsules

For the production of semi-permeable capsules, the middle phase was supplemented with 15 % porogen butyl acetate or 1-decanol (both from Sigma-Aldrich), 1 % Span80 (Fluka) surfactant and 4 % 2-Hydroxy-2-methylpropiophenone (97%, Sigma-Aldrich) photoinitiator. In the direct encapsulation of fluorophores, we supplemented the 10 % PVA inner aqueous phase with 100 *µ*g/mL of 500 kDa FITC-dextran and 50 *µ*g/mL TMR-dextran.

Fluorophore-containing capsules were washed as previously described and kept in wash buffer at 4^°^C overnight. The capsules were pipetted onto a cell-counting slide (Countess, Thermofisher) and fluorescence images were acquired in the Cy3 and FITC channels. For permeability characterization, empty capsules were immersed in a solution containing 10 kDa RITC-dextran and 32.7 kDa EGFP. After different incubation times, fluorescence images were acquired in the Cy3 and FITC channels.

### Direct encapsulation of proteins and enzymes

For direct encapsulation of proteins, the 10 % PVA inner phase was supplemented with 32.7 kDa recombinant EGFP (Biovision) mixed to a final concentration of 2 *µ*g/mL or 60 kDa FITC-labeled streptavidin (Biolegend) to a final concentration of 50 *µ*g/mL. Double-emulsions or polymerized capsules were loaded in a cell-counting chamber and imaged on a microscope with 20x magnification.

For direct encapsulation of enzymes, econoLuciferase 10 mg/mL solution (Biosynth) was mixed to 4 v/v % in the 10 % PVA inner phase. The double-emulsions and polymerized capsules were loaded in a cell-counting chamber and inspected on a microscope and fluoresence signal was measured. In the bioluminescent assay, 25 *µ*L of blank, empty capsules, econoLuciferase-containing capsules, or free econoLuciferase were mixed with 100 *µ*L of luciferase assay reagent (Promega) in a black 96-well plate and imaged for 50 minutes on a plate reader. The bioluminescent signal was integrated over the duration of the experiment. *β*-galactosidase aqueous glycerol suspension (Sigma-Aldrich) was mixed to 10 v/v % in the 10 % PVA inner phase. The produced capsules were resuspended in a 0.5 M trehalose solution and 15 *µ*L drops were pipetted onto a silicon mat and dried overnight in a 37^°^C incubator. The resulting dried pellets were placed in a tube and rehydrated with a chlorophenol red *β*-D-galactopyranoside (CPRG) solution. The color change was imaged with a mobile phone camera, or with a color camera (Pike) mounted on a microscope and imaged with a 4x objective.

### DNA-strand displacement reactions

We used capsules containing 60 kDa FITC-labeled streptavidin for the immobilization of biotinylated DNA strands. The DNA strands were synthesized by IDT with HPLC purification. The DNA sequence and DNA modifications were strictly identical to the ones presented in Joesaar *et al*.[3] and we followed the same protocol for the assembly of two populations of capsules containing the first (*F*_1_) and second (*F*_2_) gate DSD reactions. Briefly, 40 *µ*l of a dispersion of FITC-streptavidin containing capsules, 20 *µ*l of 4x buffer and 8 *µ*l of biotinylated DNA gate strand (*F*_1_ or *F*_2_, from a 10 *µ*M stock solution) were mixed with a pipette in a 1.5 ml Eppendorf tube and incubated at room temperature for 1 h, followed by addition of 12 *µ*l output strand (*Q*_1_ or *Q*_2_, from a 10 *µ*M stock solution), gentle mixing and overnight incubation at 4^°^C. The excess unbound output strand was removed by removing around 50 *µ*l of the supernatant and the capsules were washed 3 times with 400 *µ*l of buffer by centrifugation at 1500 rpm, 4^°^C, and the capsules with the immobilized DSD reaction were stored at 4^°^C. To perform the double-layer signaling cascade, the two capsules populations were mixed and supplemented with a fuel strand. After the addition of an input strand (*A*) at a concentration of 50 nM, 10 nM or 100 nM, the resulting solution was pipetted in a cell counting chamber (Countess, Thermofisher). To prevent evaporation, the device was placed in a petri dish and surrounded by kimwipes saturated with 1xPBS, allowing for long-term imaging of the two-layer signalling cascade. The fluorescent signal for the first gate (Cy5) and second gate (Cy3) was measured with a 200 ms exposure every 30s for a duration of 1 hour. The resulting images were captured in a 3×3 stitch with a 20x objective. For an estimation of the signal increase over the timecourse of the experiment, we used ImageJ and applied the “Find edges” process followed by thresholding using the default settings to detect the capsules in each image. We used erosion three times followed by dilation three times to remove smaller detected particles. Then, the process “fill holes” was used once, followed by 5 x erosion, 3 x dilation, 5x erosion, 3 x dilation, 5 x erosion. We finally used the “analyze particles” function to extract the median for particles with circularity over 0.5. For each timepoint of the experiment, the median intensity of every detected particle and the median intensity of all particles was plotted using PRISM.

## Supplementary Figures and Tables

**Figure S1:**
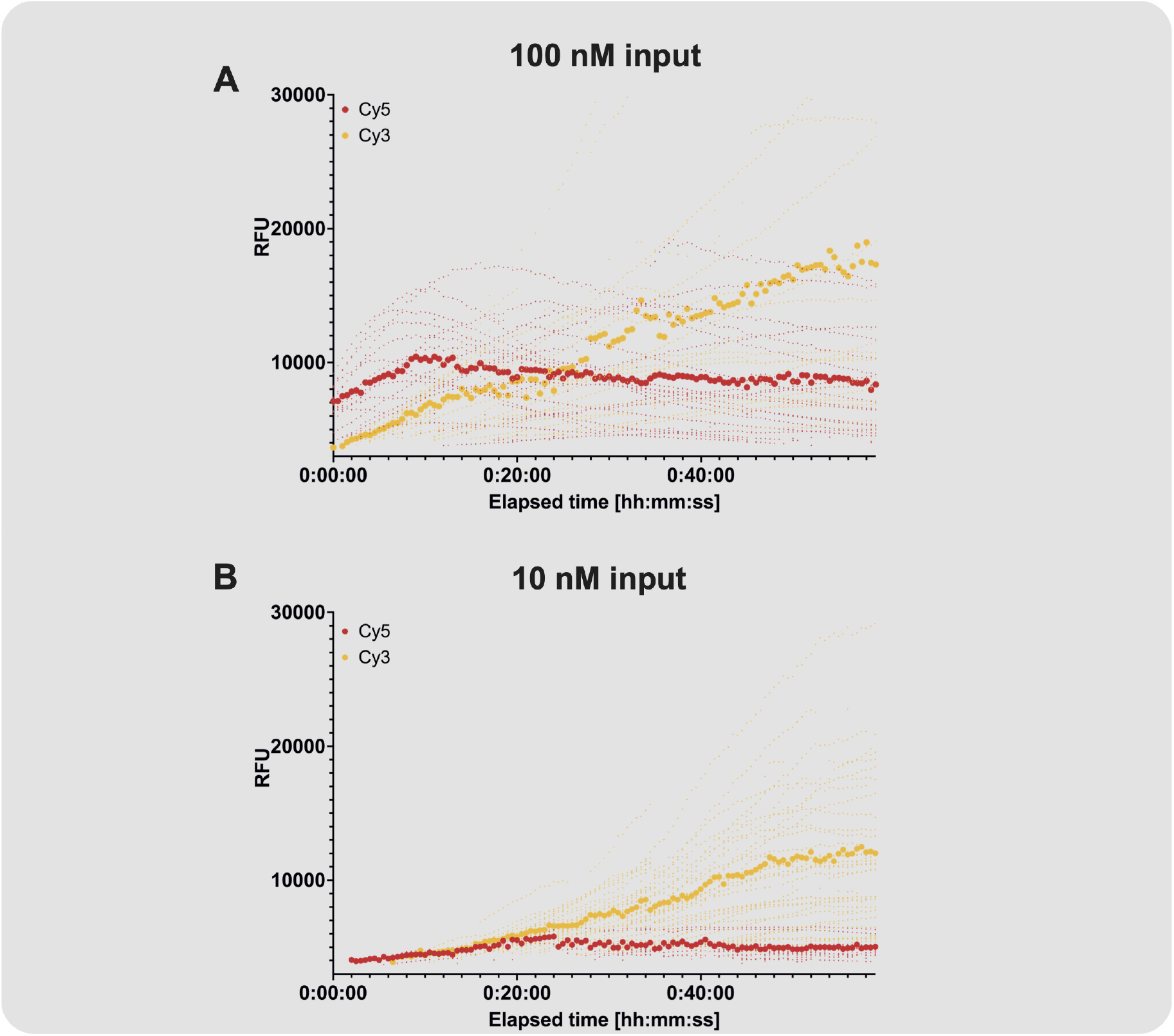
Immobilization of DNA strand displacement reaction in semi-permeable microcapsules and implementation of a two-layer signalling cascade. **(A)** Median intensity of detected particles following activation with100 nM input. **(C)** Median intensity of detected particles following activation with 100 nM input. In both plots, the larger symbols correspond to the median of all detected particles in a given fluorescent channel.

